# scREMOTE: Using multimodal single cell data to predict regulatory gene relationships and to build a computational cell reprogramming model

**DOI:** 10.1101/2021.10.11.463798

**Authors:** Andy Tran, Pengyi Yang, Jean Y.H. Yang, John T. Ormerod

## Abstract

Cell reprogramming offers a potential treatment to many diseases, by regenerating specialized somatic cells. Despite decades of research, discovering the transcription factors that promote cell reprogramming has largely been accomplished through trial and error, a time-consuming and costly method. A computational model for cell reprogramming, however, could guide the hypothesis formulation and experimental validation, to efficiently utilize time and resources. Current methods often cannot account for the heterogeneity observed in cell reprogramming, or they only make short-term predictions, without modelling the entire reprogramming process. Here, we present scREMOTE, a novel computational model for cell reprogramming that leverages single cell multiomics data, enabling a more holistic view of the regulatory mechanisms at cellular resolution. This is achieved by first identifying the regulatory potential of each transcription factor and gene to uncover regulatory relationships, then a regression model is built to estimate the effect of transcription factor perturbations. We show that scREMOTE successfully predicts the long-term effect of overexpressing two key transcription factors in hair follicle development by capturing higher-order gene regulations. Together, this demonstrates that integrating the multimodal processes governing gene regulation creates a more accurate model for cell reprogramming with significant potential to accelerate research in regenerative medicine.

## 1 Introduction

Cells generally begin their lives as a pluripotent stem cell that gradually differentiates into specialized cell fates over time. Once differentiated, cells usually have regulatory mechanisms to ensure that the cell maintains a stable state, reliably performing its required function. Recent advances in cell reprogramming have fundamentally altered our view of cell identity. Numerous experiments have established that overexpression of a few transcription factors (TFs) is sufficient to revert a differentiated cell to a pluripotent state or another specialized cell type.

These developments in cell reprogramming are significant for the field of regenerative medicine as it will facilitate the development of therapies to replenish cells our body can no longer produce. This opens the potential to regrow, repair or replace tissues and organs which may be damaged from age, disease, stress or trauma. For example, type 1 diabetes is the result of the loss of insulin-producing beta cells in the pancreas and recent experiments have shown that overexpressing the TFs *Pdx1* and *MafA* can reprogram pancreatic alpha cells into insulin-producing beta cells, effectively reversing type 1 diabetes in mice [1, 2]. Cell reprogramming has also been considered as a treatment for a wide variety of other diseases including Parkinson’s disease [3, 4], heart disease [5], spinal cord injury [6], macular degeneration [7], hearing loss [8], and aplastic anemia [9], among others.

Despite the overwhelming potential for cell reprogramming therapies to alleviate the world’s disease burden, significant roadblocks have limited our ability to perform desired cell conversions. Cell reprogramming experiments are currently very slow and inefficient, taking several weeks and producing low quantities of the desired cell type (1%; [10]). Furthermore, reprogramming is often initiated by overexpressing a combination of TFs, but there is estimated to be more than 1,500 human TFs. Many successful combinations were historically determined through a trial and error approach which is time consuming and expensive, and may not even find an optimal combination [11].

These limitations for successful cell reprogramming could be addressed with a computational model to predict the outcome of a reprogramming experiment, even to some small degree of accuracy, which could guide the hypotheses to be experimentally validated. Advances in sequencing techniques have led to the generation of large multi-omics data sets that for the first time enable a systematic view into the regulatory processes in cells. In particular, this has facilitated the development of several computational methods for cell reprogramming, taking on a range of different approaches. These include differential expression [12, 13, 14], Boolean networks [15, 16, 17], dynamical systems [18, 19], and regression [20]. However, these methods have significant limitations. Firstly, most of them assume that the cell population is homogeneous and responds to perturbations in a fixed way. However, cell reprogramming experiments have resulted in very heterogeneous outcomes with many cell subpopulations, often dependent on the initial cell state [21, 22]. Furthermore, most methods are only able to make short term predictions of the effect of TF perturbations, which may not capture the entire cell reprogramming process which involves significant changes to the cell’s identity. This motivates the need for a more holistic computational model for cell reprogramming.

Here, we present scREMOTE (single cell REprogramming MOdel Through cis-regulatory Elements), a computational method for cell reprogramming using data from simultaneous scRNA-seq and scATAC-seq. These data give us a more holistic view of the regulatory systems at the cellular level, allowing us to more accurately predict the downstream effect of the overexpression of transcription factors. We achieve this by first calculating a regulation potential, the ability of a TF to regulate a gene via cis-regulatory elements (CREs). We then build a linear regression model based on the gene expression and regulation potential, and demonstrate its applicability in predicting the effect of TF overexpression in murine hair follicle development.

## 2 Materials and Methods

### scREMOTE: a computational model to infer gene regulation and cell reprogramming

We present a novel computational model to infer gene regulation and cell reprogramming that leverages data from emerging multimodal single cell sequencing technologies. scREMOTE models four key components of gene regulation (Fig 1A) as

**Figure 1:**
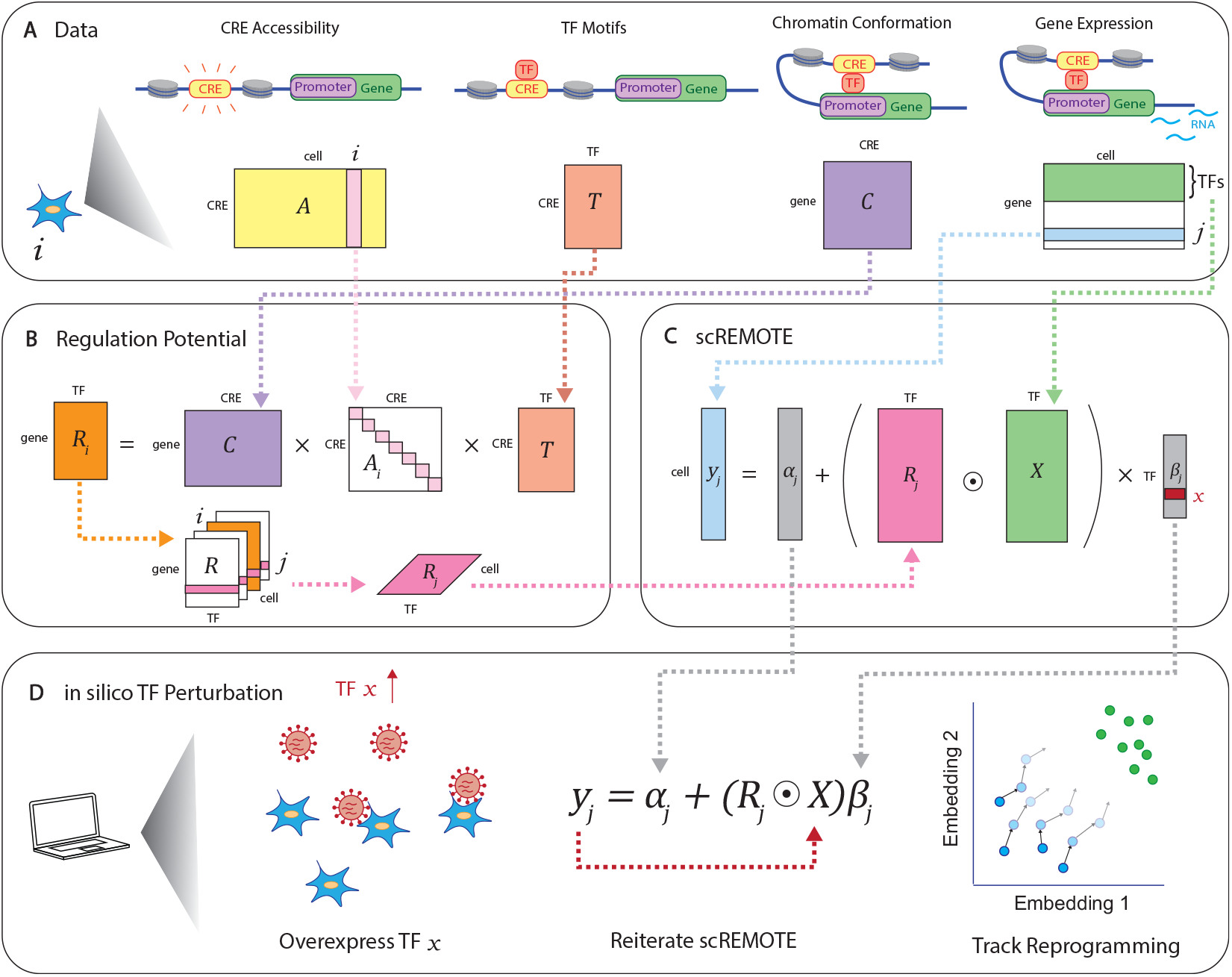
Schematic of scREMOTE. **(A)** The data inputs to scREMOTE. 1. CRE accessibility, a *CRE ×cell* matrix, 2. TF motifs, a *CRE × TF* matrix, 3. Chromatin conformation, a *gene × CRE* matrix and 4. Gene expression, a *gene × cell* matrix, where the TFs are a subset of the genes. **(B)** Calculation of binding potential. The matrix *A*_*i*_ is created by placing the CRE accessibility scores for the *i*th cell along the diagonal, and zeroes elsewhere. This has the effect of summing the regulatory potential over all CREs. **(C)** Calculation of fitted coefficients. **(D)** In silico overexpression of TF *x*.

- (1) CRE accessibility, **A**, where TFs can only bind to regions of the genome that are accessible;
- (2) TF motifs, **T**, where TFs need a matching motif in order to bind to a CRE;
- (3) Chromatin conformation, **C**, where CREs need to be able to form a DNA loop with the promoter of the target gene; and
- (4) Gene expression, **E**, which we expect to vary based on the previous three factors.

Ideally, we would want to measure all these components simultaneously in the same cell, but this is far beyond the capability of current single cell sequencing methods. Instead, we leverage on a series of recent techniques that are able to capture both the gene expression and CRE accessibility in the same cell [23, 24, 25]. Fortunately, we can reasonably assume that the motifs a TF recognises remain the same between cells. Further, chromatin conformation will be relatively stable as it is restricted by physical constraints of the 3D genome organization into topologically associated domains [26].

The first step in scREMOTE is to estimate a regulation potential by integrating data from different modalities to model the regulation of each TF onto each gene through each CRE at the single cell level. Here, the regulatory effect of a TF onto a gene via a single CRE in a cell can be interpreted as the product of the three corresponding scores in **T, A** and **C**. That is for a regulatory potential to be positive, the CRE must be (1) enriched of the TF’s motif, (2) accessible, and (3) able to form a DNA loop with the gene’s promoter. We sum up the regulatory effect from all CREs to obtain an overall measure of regulation potential of a TF to a gene in a cell (Fig 1B). It should be noted that the resulting array of regulation potentials will be rather sparse, as there are limited cases where all three conditions are met. See “Regulation potential” for more details.

The second step of scREMOTE estimates how a cell will respond to a perturbation in TF expression. We achieve this by fitting a linear regression model with the cell’s state, represented by its gene expression, as the response. We incorporate both the gene expression data and regulation potential into the predictor of our model (Fig 1C) to better incorporate the multi-level nature of gene regulation. This way, in order for a coefficient to be significant, the TF requires both regulation potential and coexpression with the gene. The advantage of a linear model is that it allows for greater interpretability, and the estimated coefficients can be used for predictions [20]. See “Model fitting and evaluation” for more details.

The final step in scREMOTE is to perturb a TF’s expression (or a combination of TF expressions), representing the process of TF overexpression, repression, or gene knock-out. We incorporate the ability for pioneer transcription factors to open up inaccessible chromatin regions. This is because they have been strongly associated with the cell fate decision making process, as it allows more TFs to bind to the DNA, further regulating gene expression [27, 28, 29]. Using the coefficients from the linear model, a change in TF expression would result in a change in the response, that is the gene expression. The predicted gene expression values of TFs can be refit into the model (Fig 1D), calculating new values for the overall gene expression, and thus cell state. This process can iterated, representing the changes over time, until convergence, resulting in the final reprogrammed state.

### Regulation potential

For a typical cell *i*, we can represent the regulatory potential, **R**_*i*_, a *gene × TF* matrix by

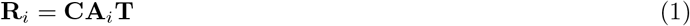

where **A**_*i*_ is a *CRE × CRE* matrix with the CRE accessibility scores for the *i*th cell along the diagonal, and zeroes elsewhere. **C** and **T** can be either binary (representing the presence) or continuous (representing the degree) of chromatin conformation and TF motif enrichment respectively. This has the effect of summing the regulatory potential over all CREs. Calculating this for all cells, we end up with a *gene × TF × cell* array which we call **R** containing the regulation potential of all transcription factors to each gene in each cell.

To verify that our calculated regulation potential is capturing true regulatory relationships, we compared our calculated values to known TF-gene regulations from the following databases: TRRUST [30], hTFtarget [31], TFBSDB [32], RegNetwork [33] and MSigDB [34, 35]. As we filtered the data to the 1000 most highly expressed genes (see Data for details), we subsetted each data set to only those in our filtered list. We also consider a Combined database, which takes the union of interactions from the 5 other databases.

As our regulation potential is at a single cell resolution, we took the average over all cells to obtain a *gene × TF* matrix to compare it to these databases. If the regulation potential is accurate, we expect that the TF-gene regulations from the databases should have a greater regulation potential than a random subset of TF-gene regulations. By resampling 1 million random subsets of the same size as each database, we compute an empirical *p*-value as the probability that the mean regulation potential from a random sample is greater than the mean regulation potential of the database interactions. This is similar to the method used by Garcia-Alonso and colleagues, where gene expression is used as a reference to benchmark TF regulation databases [36]. Here, our evaluation is in the reverse direction, using the databases as a ground truth to validate the regulation potential.

### Model fitting and evaluation

For an individual gene *j*, we propose the following model to predict its expression.

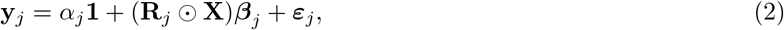

where **y**_*j*_ is the gene expression of gene *j*, **X** is the gene expression of the TFs, and **R**_*j*_ is the regulation potential corresponding to gene *j, α*_*j*_ and ***β***_*j*_ are the regression coefficients, ***ε***_*j*_ is residual noise, and represents the Hadamard product (element-wise multiplication). Note that **R**_*j*_ can be interpreted as a slice corresponding to gene *j*, with dimensions *TF × cell* of the full regulation potential matrix **R** which has dimensions *gene × TF × cell* (Fig 1B). This way, **R**_*j*_ can be interpreted as reweighting the gene expression values and the coefficients ***β***_*j*_ represent the direct effect on gene *j*’s expression when perturbing a TF.

However, we found that in practice, the **R**_*j*_ matrix was extremely sparse, which could be attributed to the fact that each of the component matrices **T, A**, and **C** are already sparse due to the nature of sequencing techniques used. This causes the coefficients in (2) to have large bias in the fitted coefficients. To alleviate this issue, we consider adding a small constant to each entry in **R**_*j*_, similar to a pseudocount or fudge factor. This has the effect that when **R**_*j*_ values are all not available (i.e., 0), for a particular TF, the regression will rely on the available gene expression values only. However, if **R**_*j*_ values are available, i.e., not all 0 for a particular TF, then they will be incorporated into the regression. This gives us a new model

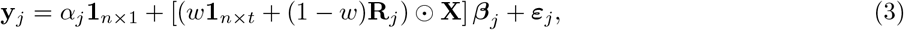

where *w* is a parameter that can be used to weight the influence of **R**_*j*_ when available. We choose to set *w* = 0.1. We saw that when *w* is small, there is minimal difference between different choices of *w*. This model was found to be the most effective when applied to experimental data, incorporating the regulation potential when available.

We compare our results to the Coexpression Model defined by:

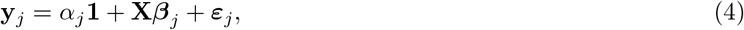

which only uses gene expression data. When fitting the model in equation 3 and 4, we use ordinary least squares regression, using the `lm()` function in `R` [37]. However, to address the situation of a TF regulating itself, that is when the response variable **y**_*j*_ is a TF, we set all the elements for the corresponding column in **X** to 0, as otherwise it would be perfectly equal to the response. We then change the fitted coefficient for the TF from 0 to 1, to encourage the TF expression level to stay constant, unless there is a perturbation.

#### In silico perturbation with scREMOTE

To predict the effect of perturbing a TF (or a combination of TFs), we can perturb the values of **X**, for example, by adding a constant to all values in the column corresponding to an overexpressed TF. The resulting changes in **y**_*j*_ represent the perturbed gene expression. To model the downstream effect of this perturbation, this process can be iterated, where the new values of the TF can be substituted into **X**, which produces a new prediction for gene expression. This iterative process can be repeated for any number of time steps or until convergence. We chose to perform our simulations for 15 time steps, as this was generally enough iterations for perturbations to converge.

To incorporate the ability for pioneer transcription factors to open up inaccessible chromatin regions, we first consider the enhancer targets of the overexpressed TF. These are defined to be the CRE’s with a score in the position weight matrix above a certain threshold, which we chose to be 0.2. We then increase the accessibility of these target CREs by adding a constant to the corresponding values in **R**_*j*_, and update the regulation potential, **R**_*j*_ in equation (3), before the perturbation.

To ensure the predicted gene expression values are biologically plausible, we can impose a minimum and/or maximum expression value. Here, any predicted expression below the minimum is replaced with the minimum value, and any predicted expression above the maximum is replaced with the maximum value. In practice, the minimum value will be 0, but the maximum would be harder to define since it would depend on the gene, cell type and sequencing depth. Some suggested ad hoc approaches for selecting a maximum could be the highest count observed in the entire data, or a few standard deviations above the mean for each gene. In our example, we imposed a minimum value of 0 and no maximum value, as predicted values seemed reasonable.

**\***Marker gene analysis. We determined gene markers by using moderated *t*-tests implemented in the `limma` package [38] in `R`. Here, we used a supervised approach, using the cell type labels provided by Ma and colleagues [23]. By performing differential expression analysis between the IRS and Hair Shaft cells, we took the top three marker genes for each cell type ranked by *p*-value, but excluded *Gata3* and *Runx1* as they will be artificially overexpressed. We should expect that a successful reprogramming model will cause the markers of the target cell type to increase in expression.

### Data

#### Chromatin conformation

The full mouse dataset was downloaded from the 4D Genome Database [39] on 21/10/2020. All coordinates were realigned from the mm9 genome to the mm10 genome using the LiftOver tool provided by the Human Genome Browser at UCSC [40]. This list of chromatin interactions is filtered down to those which include gene promoters, determined as any interactions within 500bp of the transcription start site of a gene. Gene coordinates were downloaded from the Mouse Genome Informatics (MGI) website [41].

All chromatin regions which had an interaction with a promoter were considered a CRE. These regions were sorted into bins of length 1000bp which is now taken as our CRE list. We then construct **C**, as a binary matrix indicating a recorded connection between a CRE and a gene. Usually the chromatin conformation would be measured as a score representing the strength of the connection. However, as we are using a database, the data comes from many different experiments which would not be comparable. Our final matrix **C** is a *gene × CRE* matrix.

#### TF motifs

The affinity for a TF to bind to a CRE could be estimated using TF motif enrichment or ChIP-seq data. We chose to estimate **T** using TF motif enrichment, as TF motif data is readily available on the JASPAR database [42], whereas ChIP-seq databases can often be sparse and noisy [13]. We note that only TFs present in these databases would be used in the analysis. If one wishes to include a particular TF (perhaps one that may be important for cell fate determination) which is not in the database, a baseline TF binding affinity or an estimate from ChIP-seq data may be used instead.

In our example, TF motifs were downloaded from the JASPAR database (8th release, 2020) [42] on 25/07/2020, using the full vertebrates position frequency matrices. From the CRE coordinates identified previously, the genomic sequences of our CREs were obtained using the `BSgenome.Mmusculus.UCSC.mm10` package on Bioconductor. TF motif enrichment was performed on each sequence using the AME function in the MEME Suite collection [43] with default settings. Only TFs that were highly enriched (marked as true positives) were kept, and their Position Weight Matrix score was normalized by dividing by the maximum value, so all values are between 0 and 1. This is then used as the corresponding value in **T**, a *CRE × TF* matrix.

#### scRNA-seq and scATAC-seq

The simultaneous scRNA-seq and scATAC-seq data with cell type labels from the SHARE-seq protocol was obtained from the authors upon request (S. Ma, personal communication, 23 September, 2020) [23]. This data set is now available from GEO (Accession number: GSE140203). Due to the sparsity of the gene expression data, we only used the 1000 most highly expressed genes which were then log(*x* + 1) transformed. Our gene expression matrix is a *gene × cell* matrix.

The scATAC-seq data was then realigned to match our new CRE list in 1000bp bins. As the bin cutoffs did not match exactly, any observed scATAC-seq measurement that overlapped with our 1000bp CREs was considered as a count. Applying this criteria to all our CREs gives us **A**, a *CRE × cell* matrix.

We subsetted the data to only contain the cell types of interest: Hair Shaft cells, IRS cells, and two populations of TACs which were combined. Due to a large imbalance in the numbers of each cell type, we subsampled the the larger cell types so that they all have the same size. We repeated the in-silico cell reprogramming using a range of different subsamples, and observed similar results (Fig S1, S2). All preprocessing steps and analysis were done in `R` and the code is available at https://github.com/SydneyBioX/scREMOTE.

### Visualization

PCA is the dimension reduction technique in this study for the visualization of all simulation results as we believe it is more appropriate than other sophisticated dimension reduction techniques like tSNE or UMAP commonly used in single cell research [44]. This is because PCA enables the projection of simulated data onto the same embedding as the original data, which can be used to track the changes over time. Other methods like tSNE or UMAP do not allow additional points to be projected into the embedding, unless the entire embedding is recalculated at each time point, but this would cause all cells to change positions, which cannot track the effect over time as in Fig S1-S2. As expected, we found that PC1 is strongly correlated with the total read count in each cell (Fig 2B), and it does not help to distinguish between different cell clusters (Fig 2A). However, the combination of PC2 and PC3 shows a clear separation between the TAC, IRS and Hair Shaft clusters (Fig 2C) so we use this for all visualizations.

**Figure 2:**
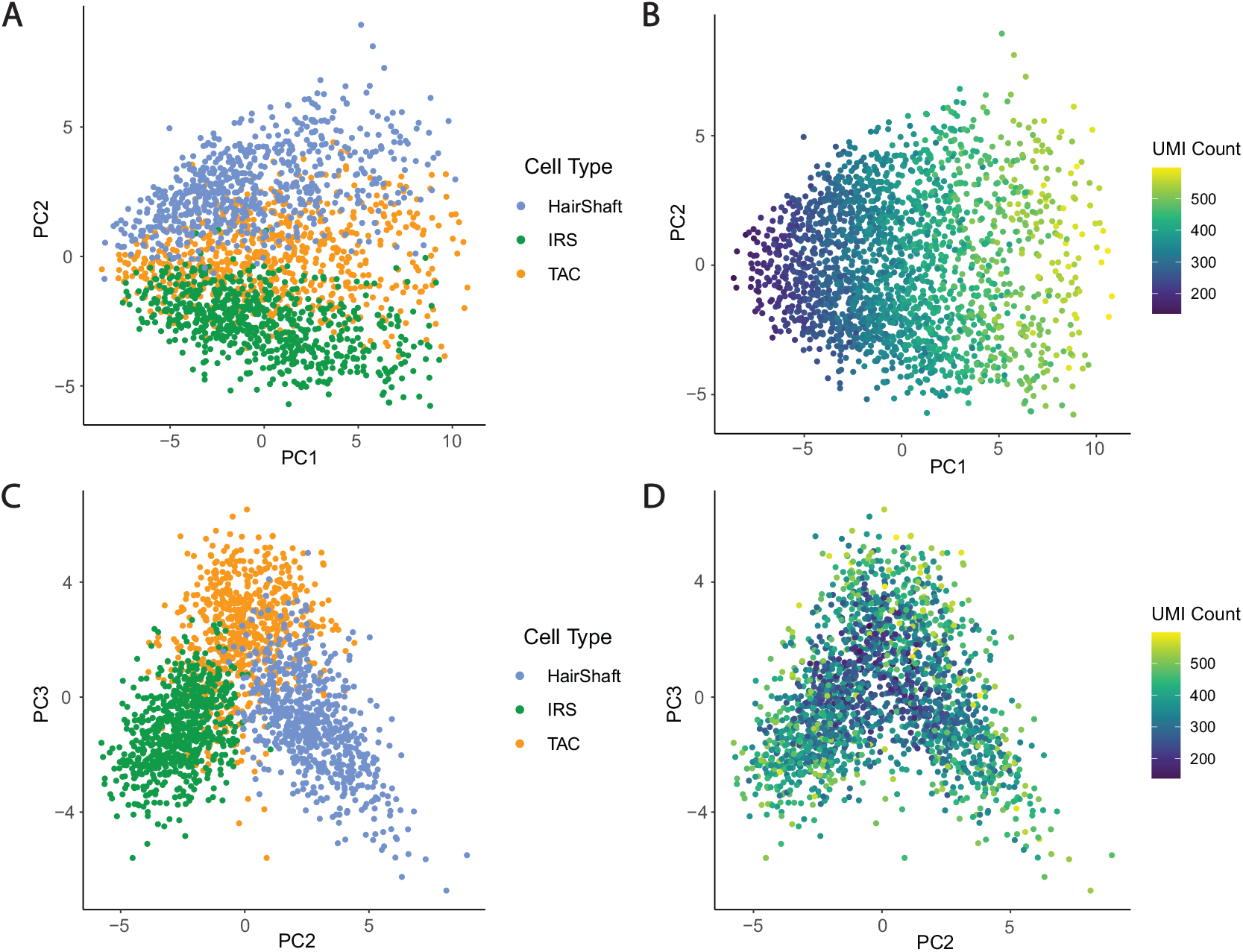
Data visualization with PCA. **(A)** Plot with PC1 and PC2, colored by cell type. **(B)** Plot with PC1 and PC2, colored by read count. **(C)** Plot with PC2 and PC3, colored by cell type. **(D)** Plot with PC2 and PC3, colored by read count.

## 3 Results

### Regulation potential captures transcription factor to gene regulations

To demonstrate the applicability of scREMOTE, we investigate the hair follicle developmental system, as it is a natural differentiation system in the adult skin and has been profiled with simultaneous scRNA-seq and scATAC-seq [23]. Furthermore, the ability to reprogram hair cells in the inner ear could be used as a cure to permanent hearing loss [45]. We chose to estimate **T** using TF motif enrichment and we estimated **C** using data from the 4D Genome Database [39]. See *Materials and Methods* for more details.

We first verify that our calculated regulation potential is capturing true regulatory relationships, by comparing our calculated values to known TF-gene regulations. There are a variety of databases that record known and predicted regulations, such as TRRUST [30], hTFtarget [31], TFBSDB [32], RegNetwork [33] and MSigDB [34, 35]. As expected, we find that the TF-gene regulations from these databases have a greater regulation potential than random subsets of TF-gene pairs. By resampling 1 million random subsets for each database, we see that all databases are significantly enriched with interactions containing a high regulation potential, implying that our regulation potential captures true regulatory TF-gene relationships (Table 3).

### scREMOTE predicts the outcome of cell reprogramming experiments

We now show how scREMOTE can be used to perform an in silico TF overexpression experiment. In the hair follicle developmental system, Transit-Amplifying Cells (TACs) differentiate into either the Inner Root Sheath (IRS) or Hair Shaft lineages. *Gata3* has long been identified as a reprogramming TF for the IRS lineage [46, 47] and also *Runx1* for the Hair Shaft lineage [48, 49, 50]. Thus, we expect that an accurate model will predict that an overexpression of *Gata3* will reprogram the TACs towards the IRS cells (Fig 3A), and that an overexpression of *Runx1* will reprogram the TACs towards the Hair Shaft cells (Fig 3B). In the case of unsuccessful reprogramming, we would see the perturbed cells converge to a different cluster. To demonstrate the value of multimodal data in scREMOTE, we compare it to an equivalent model which only uses gene expression, which we call the Coexpression Model, as it would only detect linear coexpression patterns.

**Table 1:** Results of testing regulation potential with TF-gene regulation databases. The empirical *p*-value is calculated to be the probability that the mean regulation potential from a random sample (of the same size as the database) of TF-gene pairs is greater than those from the database.

| Database | Number of Regulations | Empirical p-value |
| --- | --- | --- |
| TRRUST | 43 | 0.03 |
| hTFtarget | 10294 | $< 1 \times 10^{-6}$ |
| TFBSDB | 5253 | $< 1 \times 10^{-6}$ |
| RegNetwork | 729 | $3.9 \times 10^{-5}$ |
| MSigDB | 695 | $< 1 \times 10^{-6}$ |
| Combined | 14112 | $< 1 \times 10^{-6}$ |

**Figure 3:**
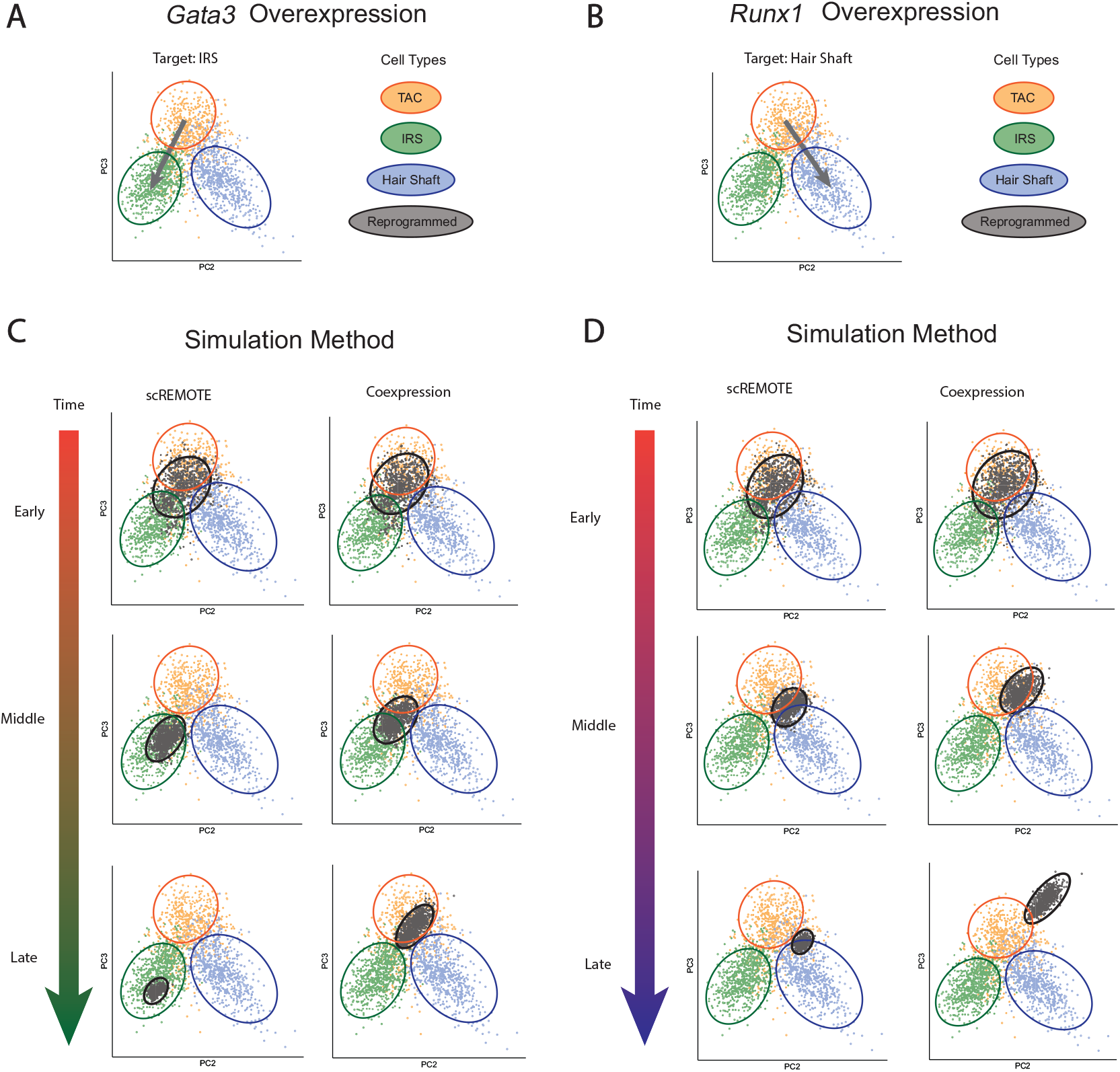
Perturbation of TFs in hair follicle development. **(A)** Expected result of *Gata3* overexpression. **(B)** Expected result of *Runx1* overexpression. **(C)** Simulated result of *Gata3* overexpression with both scREMOTE and Coexpression Model. **(D)** Simulated result of *Runx1* overexpression with both scREMOTE and Coexpression Model.

Fig 3 reveals the prediction of both scREMOTE and the Coexpression Model for overexpressing *Gata3* (Fig 3C) and *Runx1* (Fig 3D) in the TACs. We can see that when *Gata3* is overexpressed, both scREMOTE and the Coexpression Model make accurate short term predictions (early and middle time points), perturbing the cells towards the IRS cell fate. However, we see that only scREMOTE produces an accurate long term prediction (late time point). Likewise, when *Runx1* is overexpressed, we can see that both models make accurate short term predictions, perturbing the cells towards the Hair Shaft cell fate. But again, we see that only scREMOTE produces an accurate long term prediction. We believe that this is because scREMOTE is, to some extent, capturing the regulatory dynamics driving cell reprogramming whereas the Coexpression Model is limited by the highly correlated nature of gene expression. Animations of the entire reprogramming process can be found in Fig S1-S2. We see that in most cases, reprogrammed cells converge to the expected cell cluster, however occasionally a distinct cluster is formed. We believe that this is because scREMOTE has not captured the entire regulatory dynamics, involving undetected TFs and genes, and other factors like microRNAs and DNA methylation. This means that the predicted cell cluster may not match the true cell cluster as seen in the data.

### Computationally reprogrammed cells show markers of cell identity

To verify the fidelity of the scREMOTE predicted cell state to the true target cell state, we tracked the expression of several marker genes over time. Here, we should expect that successful reprogramming will have the markers of the target cell type increase in expression and the markers of the opposing cell type to potentially decrease in expression. In the overexpression of *Gata3* (Fig 4A), we see that the IRS markers all increase in expression but the hair shaft marker *Lef1* decreases, and the other hair shaft markers *Trps1* and *Kcnh1* increase to a small extent. Similarly, in the overexpression of *Runx1* (Fig 4B), we see that all Hair Shaft markers increase in expression and all IRS markers decrease in expression. This demonstrates that the predicted cell state from scREMOTE has an elevated expression of marker genes, verifying the validity of the simulated cell reprogramming.

**Figure 4:**
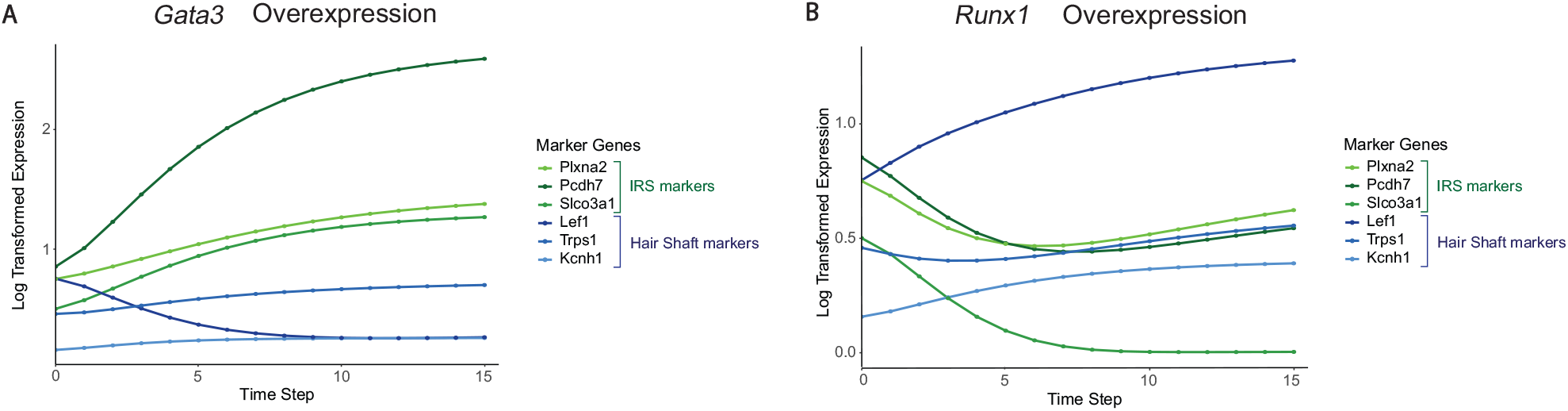
Tracking marker genes. **(A)** Marker genes during *Gata3* overexpression. Genes colored in green represent IRS markers and genes colored in blue represent Hair Shaft markers. **(B)** Marker genes during *Gata3* overexpression.

Interestingly, in the *Runx1* overexpression, we observe a reversal in the expression of the IRS markers *Plxna2* and *Pcdh7* which decrease and then increase, and also for the Hair Shaft marker *Trps1* which decreases and then increases. This suggests that scREMOTE is able to capture, to some extent, higher order gene regulations where the downstream targets of *Runx1* are causing a reversal in the trend, allowing scREMOTE to make accurate long term predictions. In contrast, the Coexpression Model is only able to predict the immediate effects of the perturbation, which may not capture the downstream effects that result in successful cell conversion.

## 4 Discussion

Understanding the regulatory dynamics in a cell is a complex yet important challenge, especially in the context of cell reprogramming, which results in large changes to the cell’s identity. Here, we present scREMOTE, a model for long-term predictions of TF perturbations at the single cell level that extends on existing algorithms [20]. Integrating simultaneous scRNA-seq and scATAC-seq data provides a more comprehensive view of the regulatory dynamics occurring in each cell. By aggregating the regulatory effect through each CRE, a regulation potential is calculated between each TF and gene. We then combined the regulatory potential and TF expression to construct a linear model for gene expression. By iteratively updating gene expression, we are able to predict the long term effects of TF perturbation. We demonstrated scREMOTE on experimental data, revealing that it can successfully model the cell reprogramming process, and capture higher order levels of gene regulation.

scREMOTE, like any computational model, is based on a set of assumptions that, to varying degrees, reflect the underlying biology. Here, we approximate the regulatory effect of each TF to be linear and additive, whereas in reality, TFs often work in combinations and in complex relationships [51]. Although, in our current implementation, we used ordinary least squares linear regression, the scREMOTE framework could be easily extended to more advanced models, possibly capturing non-linear effects or it could be extended to include regularization like LASSO if there are too many TFs, or if multicollinearity is a concern. The choice of these extended models will be motivated by the specific data structure. However, in our illustration, we found that after filtering, only 35 TFs were considered highly expressed. These TFs were not strongly correlated with each other (Fig S3), and thus regularization is unlikely to have a significant impact for our data.

Multimodal single cell sequencing technologies are still in their infancy so there is very limited data to evaluate scREMOTE under a practical setting. For our validation, we required a source cell type that could be reprogrammed into at least one (but ideally more) target cell types which show clear separation when visualized with PCA, where the reprogramming is driven by different key TFs that are sufficiently expressed. Despite this limitation of our ability to validate the scREMOTE workflow more widely, we believe that the currently available data gives us a convincing example to support the applicability of scREMOTE. In particular, it will be challenging to evaluate the performance of scREMOTE in its ability to model lowly expressed TFs, which may be very important for cell reprogramming. This challenge is due to the bias in the estimated coefficients as a consequence of most high throughput sequencing technologies which leads to a high proportion of dropouts [52]. To date, the single cell research community has employed a variety of imputation methods to provide a partial solution to this issue, but there is no consensus on the most appropriate or optimal approach [53]. Going forward, with the constant improvement in these sequencing technologies and their increasing accessibility [54, 24], we expect that in the future, cell reprogramming predictions from scREMOTE will become more applicable with increased accuracy and availability of such data.

The chromatin conformation data would ideally be measured in the same cells as the other modalities, however this is not feasible with current sequencing protocols. Also, chromosome conformation capture techniques like Hi-C are currently expensive to run with multiple complex experimental steps [55], and so it is difficult to obtain this type of data. Further complicating this component is that Hi-C has very low resolution (up to 10kb) [56] making it difficult to use for the precision required to model CRE-gene interactions. We bypassed these issues by using a database of measured interactions [39] from a range of chromatin conformation techniques which we used as a baseline measure for the chromatin conformation in all cells. As chromatin capture technologies improve and become cheaper, we will be able to collect more Hi-C data, extending the applicability of scREMOTE.

The TF motifs data is dependent on publicly available databases which are currently incomplete, for example the JASPAR database currently contains 592 profiles for mouse TFs out of an estimated 1640 [42]. This limits the applicability of scREMOTE to model TFs which may be important for cell fate determination but whose binding profile has not yet been characterized. However, with the regular update of these databases [42], scREMOTE will have continued expansion of the number of TFs that could be incorporated.

In summary, our method is the first to our knowledge that simulates cell reprogramming experiments by modeling gene regulatory systems at the single cell level through the integration of matched scRNA-seq and scATAC-seq data. The ability of scREMOTE to model the biological mechanisms behind cell reprogramming at the single cell level would lead to its increased applicability over earlier methods. We hope that this will contribute to our understanding of the role of gene regulation in cell identity and accelerate research in regenerative medicine by predicting the key TFs for new cell conversions.

## Data and code availability

All code written in support of this publication is publicly available at https://github.com/SydneyBioX/scREMOTE

## 5 Supplementary material

**S1 Fig. *Gata3* overexpression for different subsamples**.

**S2 Fig. *Runx1* overexpression for different subsamples**.

**S3 Fig. Correlation matrix of TF expression**.

## Acknowledgements

We thank all of our colleagues at the Sydney Precision Bionformatics Alliance for their support and intellectual engagement.

