## Supplementary Figure 1 for "scREMOTE: Using multimodal single cell data to predict regulatory gene relationships and to build a computational cell reprogramming model"

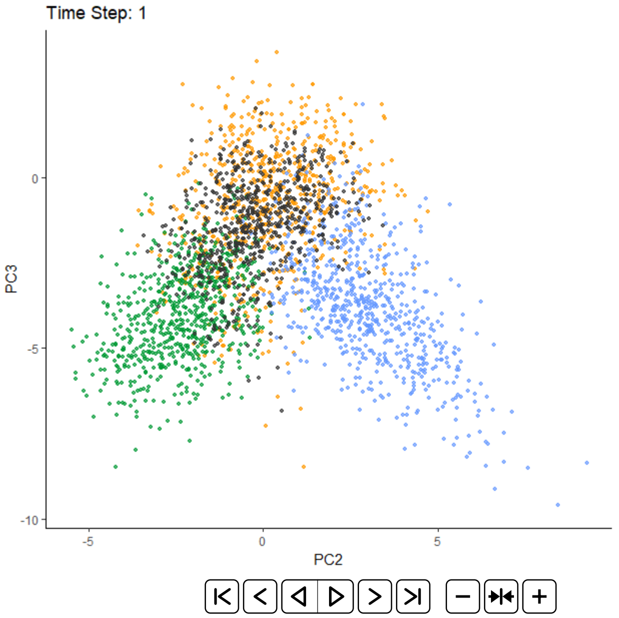


(a) Seed: 2021

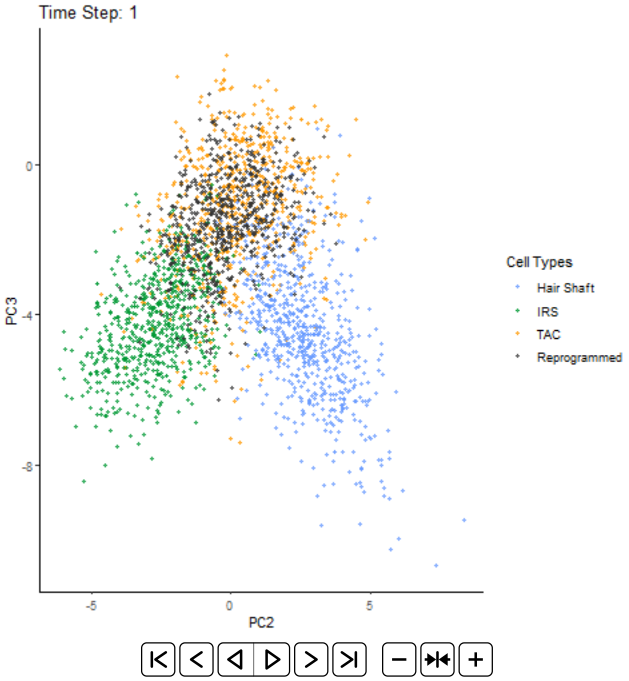


(b) Seed: 2020

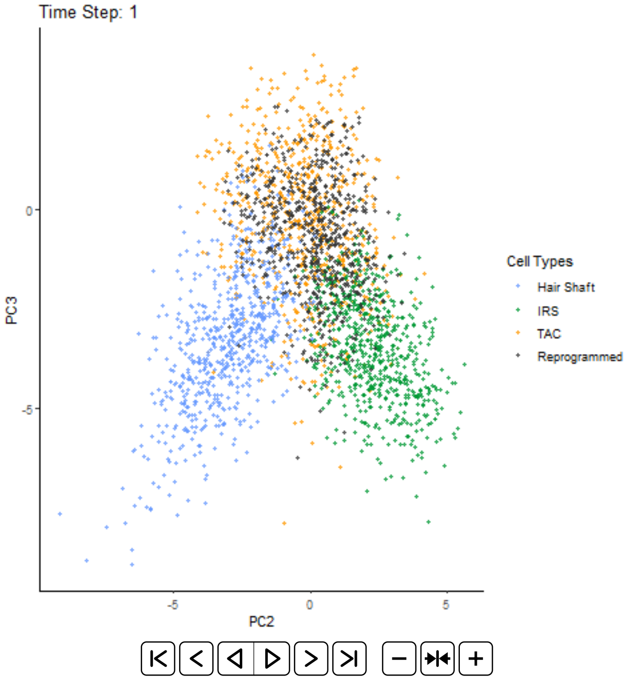


(c) Seed: 2019

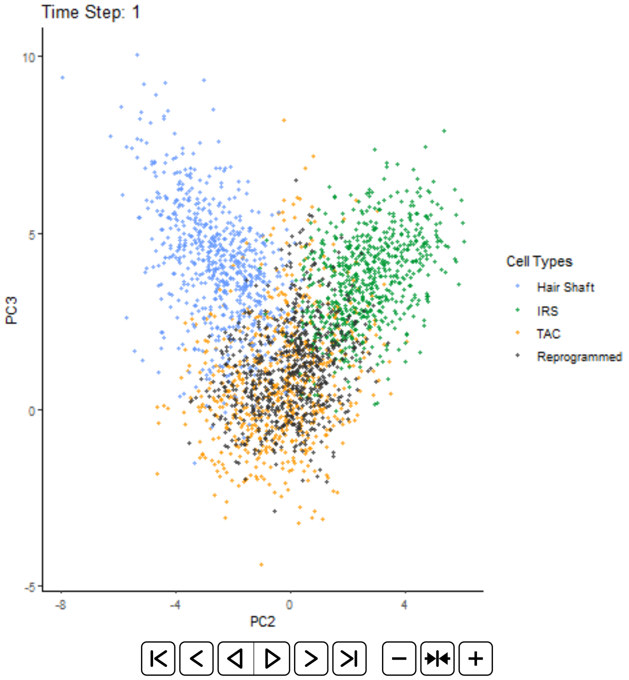


(d) Seed: 2018

Figure S1. Simulated *Gata3* overexpression with scREMOTE using different seeds
