## Supplementary Figure 2 for "scREMOTE: Using multimodal single cell data to predict regulatory gene relationships and to build a computational cell reprogramming model"

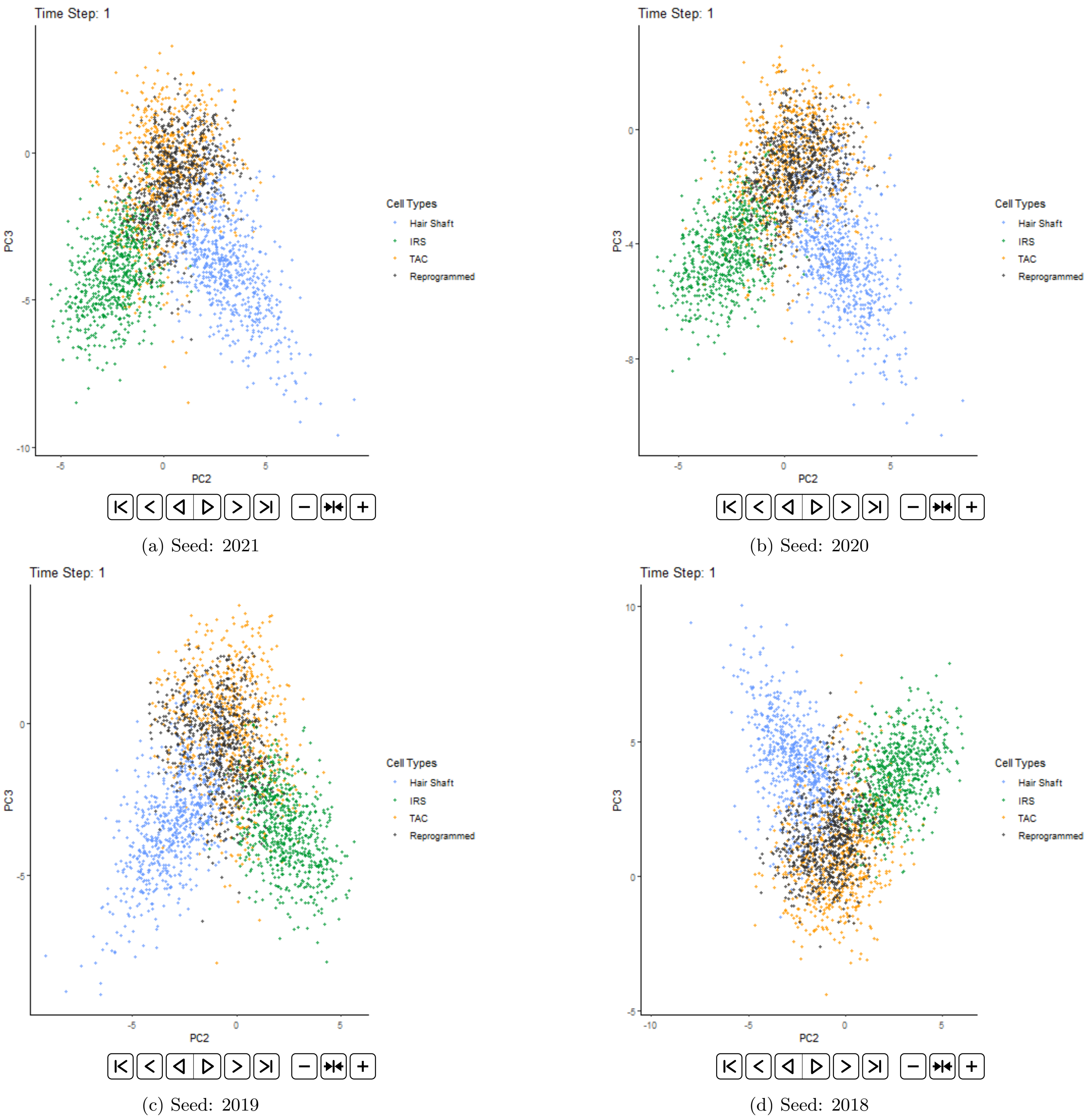


(a) Seed: 2021

(b) Seed: 2020

(c) Seed: 2019

(d) Seed: 2018

Figure S2. Simulated *Runx1* overexpression with scREMOTE using different seeds
