## Supplementary figures and images for "scREMOTE: Using multimodal single cell data to predict regulatory gene relationships and to build a computational cell reprogramming model"

### Supplementary Figure 3

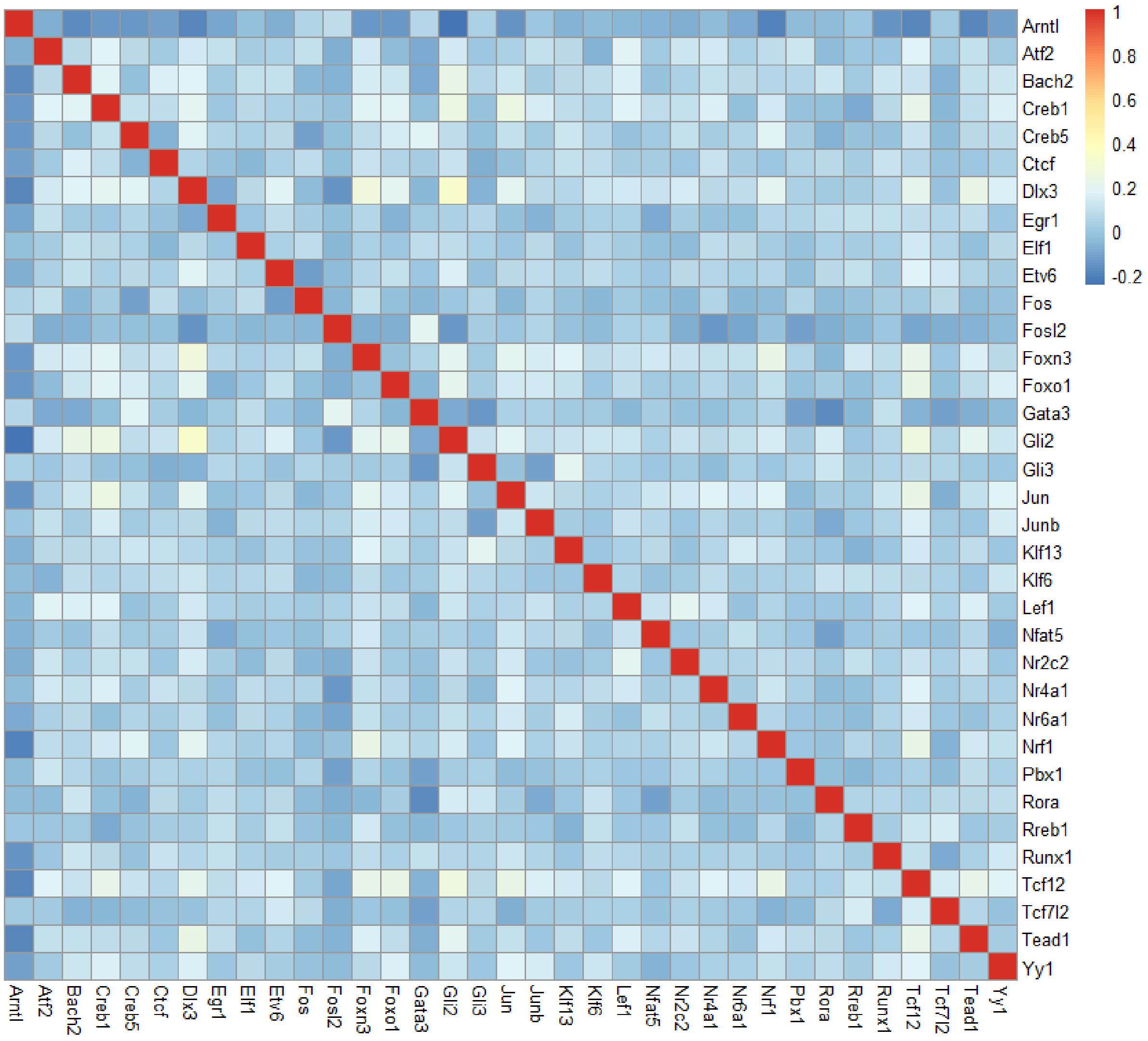


Figure S3. Correlation matrix of TF expression.
